# A modular toolbox for *in cellulo* screening of small molecule inhibitors targeting chromatin reader domains

**DOI:** 10.1101/2025.09.06.674632

**Authors:** Davide C. Recchia, Richard Cardoso da Silva, Niklas Kupfer, Angela Topic, Sara Giuliani, Rodrigo Villaseñor, Tuncay Baubec

## Abstract

The dysregulation of bromodomain proteins, a family of “reader” proteins that recognize the critical post-translational modification of acylation, is implicated in diseases like cancer, making them important therapeutic targets. However, the development of specific small-molecule inhibitors is hindered by the lack of robust, high-throughput cellular assays to measure target engagement and off-target binding in living cells. To address this gap, we developed a modular platform of cell lines that stably express synthetic chromatin reader constructs, termed Acyl-eCRs, containing various bromodomains fused to eGFP. We demonstrate that these Acyl-eCRs recapitulate the same response to bromodomain inhibitors and PROTACs as endogenous proteins, allowing for the quantitative assessment of drug effects. We introduce two complementary flow cytometry-based assays to evaluate inhibitor-target engagement: a competitive binding assay leveraging PROTAC-induced degradation, and a nuclear retention assay that directly measures the displacement of bromodomains from chromatin. Our approach circumvents the need for laborious protein purification and in vitro characterization, providing a scalable and physiologically relevant method for assessing inhibitor potency and specificity. This platform represents a versatile tool for chemical biology, enabling the functional evaluation of chromatin-targeting drugs in a native cellular context.

## Introduction

Post-translational modifications (PTMs) play an important and diverse role in regulating cellular processes. In particular, acylation, the covalent attachment of an acyl group to a molecule, stands as a profoundly prevalent and critical PTM across diverse biological systems. Its ubiquity is particularly evident on histones, where it fundamentally impacts chromatin structure and gene regulation, but it is also a widespread modification on numerous non-histone proteins, influencing their function, localization, and interactions. This versatile modification is intricately involved in a myriad of fundamental biological processes, ranging from transcription and translation to DNA replication, metabolism, and signal transduction. This PTM can also range in a variety of chemical moieties that can each add unique functional implications^1–3^, which will collectively be referred to as “acylation” from here on.

The dynamic nature of protein acylation is governed by a sophisticated system of dedicated proteins: writers, the enzymes that add acyl groups; readers, the protein domains that recognize and bind to acylated sites; and erasers, the enzymes that remove acyl groups, all of which collectively control the cellular acylation landscape^2^. Among the key reader proteins that specifically recognize acetylated (and sometimes other acylated) lysine residues are three major families of protein domains: Bromodomains (BRDs), Double Plant Homeodomain (DPF) domains, and YEATS domains^4^. Proteins containing these acyl-reader domains are frequently found to be mutated or dysregulated in various human cancers and other diseases, underscoring their crucial roles in cellular homeostasis and pathology. Consequently, these domains and the proteins they reside within represent promising targets for the development of therapeutic strategies^5,6^.

Much effort has been invested in generating drugs to target these acylation-related diseases, resulting in several drugs that show promise in both preclinical and clinical settings. These drugs can be classified as small-molecule inhibitors that compete with the endogenous targets for the bromodomain binding pocket, or as proteolysis targeting chimeras (PROTACs) that specifically degrade their protein targets^7^. These bromodomain inhibitors, especially those against the Bromodomain and ExtraTerminal (BET) family^8^, have shown promise in preclinical and clinical settings. Despite this progress, developing effective inhibitors of bromodomains has remained a significant challenge due to the shallow and often solvent-exposed nature of their binding pockets, which makes achieving high potency and selectivity difficult^9,10^. A critical hurdle in their development is the precise measurement of off-target binding within living cells, as there is currently no straightforward, direct readout to comprehensively assess such interactions. This stands in contrast to modalities like PROTACs, where the effects of target engagement and degradation can be more readily tracked by measuring protein abundances by mass spectrometry, revealing specific and unspecific targets. Consequently, the specificity of these inhibitors is often primarily determined by *in vitro* assays, such as Isothermal Titration Calorimetry (ITC) or by bespoke bromodomain engagement assays^11,12^. While valuable, these *in vitro* methods fail to fully capture the intricate cellular context, including protein-protein interactions, PTMs, subcellular localization, and competition for endogenous ligands. Furthermore, the generation of comprehensive toolboxes for these studies, as exemplified by efforts to characterize the bromodomain family, often necessitates the laborious purification of numerous individual proteins for biophysical characterization, representing a significant bottleneck^11^. Thus, a major missing component in the field is a robust, cell-based method to accurately measure off-target inhibitor binding and validate their specificity for a large number of bromodomain proteins in a physiologically relevant environment.

To address this technology gap, we develop a panel of cell lines expressing synthetic bromodomains from various proteins and demonstrate how they can be used to measure inhibitor specificity *in cellulo*. Based on our previous work, we built upon the engineered Chromatin Reader (eCR)^13,14^ concept to generate 20 cell lines expressing unique bromodomain constructs representing all families of the bromodomain family tree. These bromodomain constructs, hereafter termed Acyl-eCRs, take bromodomains in several valencies and fuse them to GFP for readout via flow cytometry & fluorescence microscopy. We demonstrate that Acyl-eCRs respond to drug perturbations in the same manner as their full-length endogenous protein counterparts and that Acyl-eCRs proved highly sensitive to drug perturbations, allowing for the quantification of inhibitor interactions across different bromodomain types, valencies, and combinations. Lastly, we show how Acyl-eCRs can be used to measure the *in cellulo* specificity of an inhibitor via competitive binding assays with PROTACs and through flow cytometry-based nuclear retention assays.

## Results

### Acyl-eCRs recapitulate the effects of drug perturbations on endogenous targets

To establish an *in cellulo* platform for the systematic analysis of bromodomain inhibitors and PROTACs, we employed engineered chromatin readers (eCRs)^13^. These eCRs are stably expressed in mouse embryonic stem cells (mESCs) and are fused in frame to a green fluorescent protein (eGFP), enabling quantification by microscopy and flow cytometry (Figure 1a). To generate Acyl-eCRs, we cloned the cDNA sequences of the bromodomains from the homologous histone acetyltransferases CBP or p300, fused in tandem to a nuclear localization signal (NLS) and eGFP into the expression cassette downstream of a CAG promoter for constitutive expression. These constructs were stably integrated into a defined locus in the mouse genome via recombinase-mediated cassette exchange (RMCE), allowing rapid generation of mESC lines expressing the eCRs of interest. This controlled genomic context ensures that all eCRs are expressed from the same promoter and locus, thereby enabling direct comparative analysis (Supplementary Figure 1a). To assess whether these Acyl-eCRs accurately recapitulate the response of the endogenous CBP or p300 to bromodomain inhibition or targeted degradation, we furthermore generated mESC lines in which the endogenous *Crebbp* or *Ep300* genes were C-terminally tagged with eGFP using CRISPR-Cas9 (Figure 1a, Supplementary Figure 1b). Successful homozygous tagging was confirmed by PCR followed by Sanger sequencing (Supplementary Figure 1c-d). Flow cytometry analysis further confirmed that these tagged proteins are stably expressed in the entire population (Supplementary Figure 1e).

**Figure 1.**
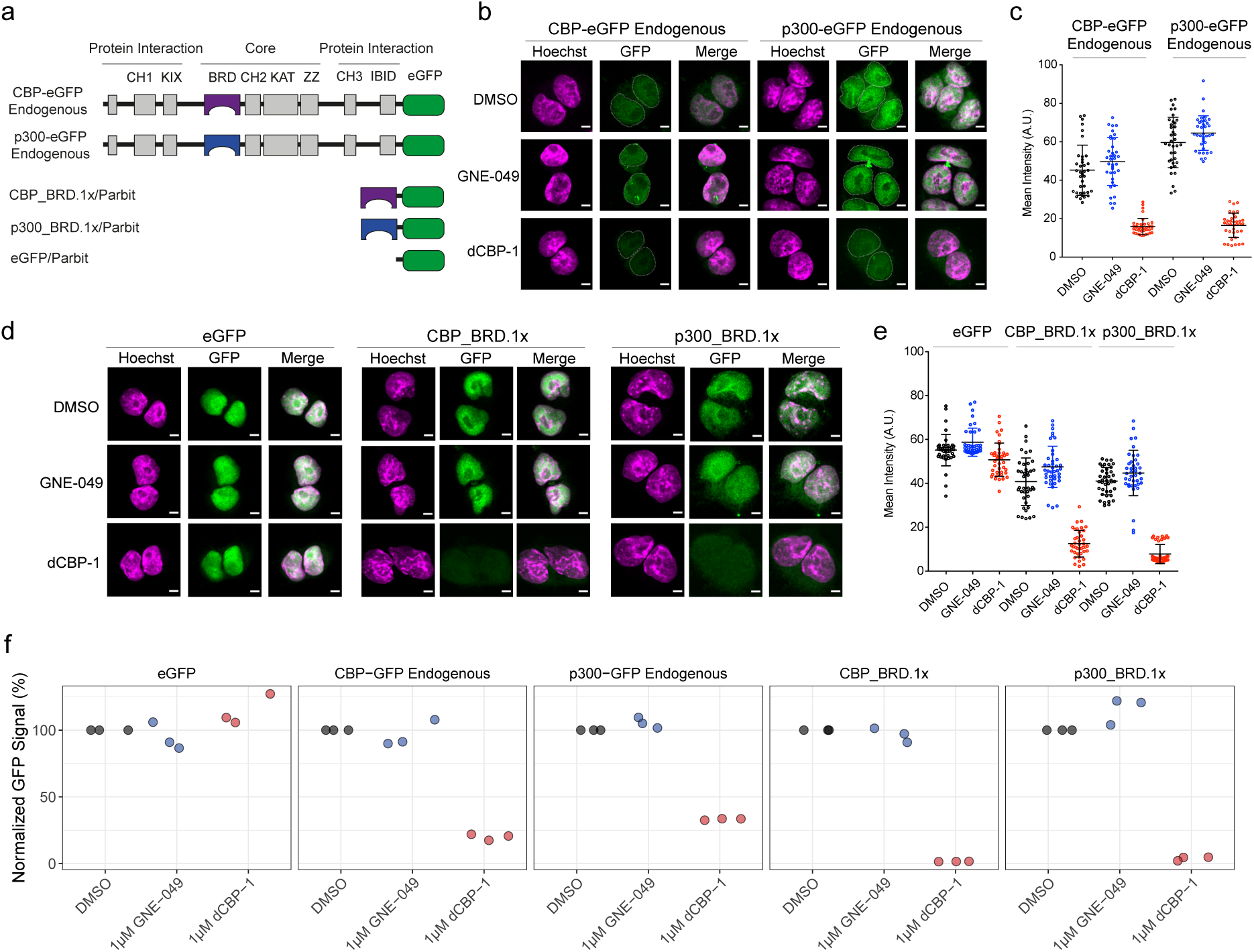
**(A)** Schematic showing how the bromodomains of CBP and p300 are being used as chromatin readers in the Parbit system, relative to their full-length endogenous proteins. **(B-E)** Immunofluorescence analysis of mESCs showing nuclear localization of CBP and p300 constructs after drug treatment. **(B, D)** Representative images of mESCs expressing either endogenous CBP-eGFP or p300-eGFP fusions **(B)**, or synthetic CBP/p300 Acyl-eCR constructs (CBP_BRD.1x/Parbit, p300_BRD.1x/Parbit, or eGFP/Parbit) **(D)**, with GFP signal in green and nuclear Hoechst staining in magenta. Cells were treated with DMSO, GNE-049, or dCBP-1 at 1 µM for 24 hours. Scale bars: 5 µm. **(C, E)** Quantification of mean nuclear GFP intensity (arbitrary units) for endogenous fusions and Acyl-eCR constructs **(E)**. Data represent mean ± SD from 40 cells per condition. **(F)** Normalized FACS data showing the effects of drug treatments on cells either expressing endogenous proteins fused to eGFP or Acyl-eCR constructs. The percentage represents the GFP signal in treated cells as a ratio of the signal observed in untreated samples of the same cell type, after normalizing for drug-induced autofluorescence in wild-type cells. Drug treatments were performed for 24 hours.

We next compared how the Acyl-eCRs and the endogenous CBP and p300 respond to inhibitor treatments specific for the CBP/p300 bromodomains. Towards this, we used the highly specific bromodomain inhibitor GNE-049, which outcompetes endogenously acylated proteins for binding to the bromodomains of CBP and p300^15,16^. In addition, we used dCBP-1, a PROTAC that selectively degrades CBP/p300 by targeting their homologous bromodomains^17^. Fluorescence microscopy using these cell lines following GNE-049 treatment shows no effect on GFP levels, suggesting that blocking the bromodomain does not directly affect protein stability (Figure 1b-e). We observed that the GNE-049 treatments cause foci formation in the endogenously tagged CBP and p300 cell lines (Figure 1b), whereas no foci formation is observed in the Acyl-eCR cell lines (Figure 1d). This is expected, since the foci formation of the full-length proteins is known to be caused by interactions with transcription factors via intrinsically disordered protein domains of CBP/p300, which are not included in the Acyl-eCR constructs^18^. In contrast, treatment with the dCBP-1 PROTAC caused the degradation of the endogenous proteins as well as the Acyl-eCRs (Figure 1b-e). The quantification of these images shows that the Acyl-eCRs are even more sensitive to the PROTAC-induced degradation than the endogenous proteins (Figure 1c and e). Western blot analysis further confirmed that the dCBP-1 PROTAC can efficiently degrade Acyl-eCRs (Supplementary Figure 1f).

To account for differences in autofluorescence resulting from the different drug treatments, and to further establish a setup that allows quantitative and high-throughput measurements, flow cytometry analysis was performed. Fluorescent signals from the eGFP-tagged proteins were obtained by gating for living and single-cell populations of mESCs (Supplementary Figure 2a), which gave robust and reproducible signals (Supplementary Figure 2b). Fluorescence signals were normalized to each treatment’s background signal to account for autofluoresence from each drug treatment. This again confirmed that the CBP and p300 Acyl-eCRs are degraded by the dCBP-1 PROTAC more efficiently than the full-length endogenous proteins (Figure 1f).

To obtain a better view of the suitability of the Acyl-eCRs for testing small molecule inhibitors, we performed time-course experiments using the dCBP-1 PROTAC on the Acyl-eCR and endogenously tagged cell lines. We observed that the Acyl-eCRs were degraded faster and more efficiently than their endogenous protein counterparts, indicating their sensitivity towards dCBP-1 (Supplementary Figure 2c). Dose-response treatments of the dCBP-1 PROTAC using the CBP and p300 cell lines showed again that the Acyl-eCRs are highly sensitive to drug perturbations and were efficiently degraded (Supplementary Figure 2d). The observed degradation in the absence of the full-length protein context suggested that these CBP/p300 Acyl-eCRs are suitable tools for drug screening *in cellulo*.

### Acyl-eCRs provide a modular platform to functionally test inhibitor specificity

We next investigated whether other PROTACs retain their ability to degrade their target bromodomains in the absence of the full-length protein context. To test this, we focused on the bromodomains of BRD4 using BRD4-specific Acyl-eCR cell lines in the presence of two compounds: the BRD4-specific PROTAC ARV-825^19,20^ and the BRD4-specific bromodomain inhibitor JQ1^8^. Additionally, we sought to determine how bromodomain valency influences PROTAC-induced degradation. To investigate this, we generated Acyl-eCR constructs containing different numbers and types of BRD4 bromodomains. This also included point mutants of both bromodomains from BRD4, which are known to abrogate binding of the bromodomain without causing structural changes^21^ (Figure 2a).

**Figure 2.**
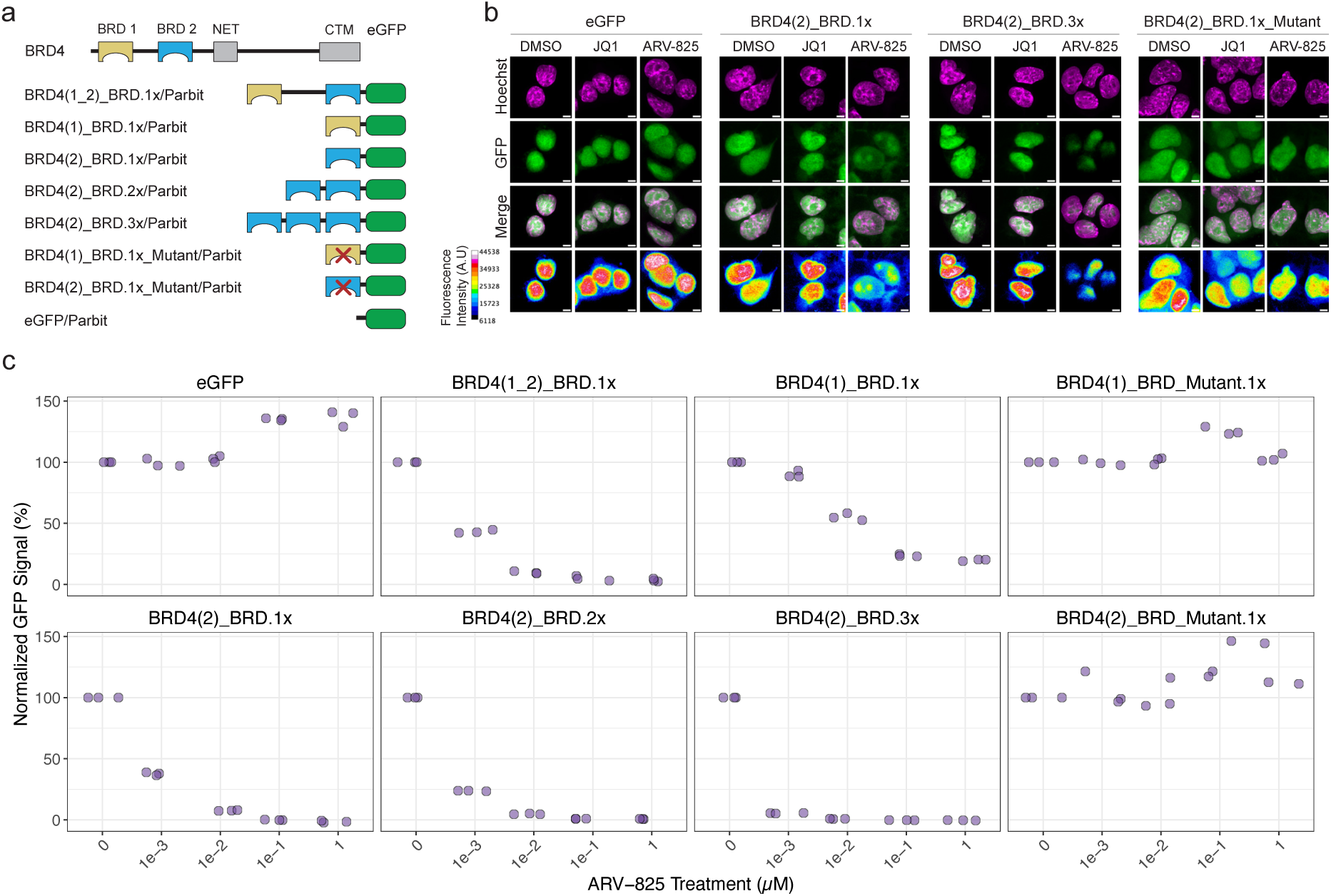
**(A)** Schematic showing the domain architecture of the Brd4 protein, and how its bromodomains are being used in several combinations to make acyl-eCRs and to determine how the valency of reader domains affects drug perturbations. **(B)** Immunofluorescence images of mESCs showing the nuclear localization of different valencies of the second bromodomain from BRD4 in the Parbit system (green) and their colocalization with Hoechst (magenta) after drug treatments. All scale bars are 5 µM. Drug treatments were performed at 1 μM concentrations for 24 hours. Bottom panel: Representative pseudocolored images (eGFP signal) depicting the differences in fluorescence intensities in different cell lines. A gradient pseudocolor bar (signal intensity) is shown at the left. **(C)** Normalized FACS data showing the effects of ARV-825 PROTAC treatment on cells expressing several combinations of bromodomains from BRD4. The percentage represents the GFP signal in treated cells as a ratio of the signal observed in untreated samples of the same cell type, after normalizing for the autofluorescence of the drug treatment in wild-type cells.

Fluorescence microscopy showed that the BRD4 PROTAC, ARV-825, is also capable of degrading its intended bromodomain targets in the absence of the full protein context. This is seen with the efficient degradation of Acyl-eCRs that contain 1x or 3x copies of the second bromodomain from BRD4. It is also shown that the degradation of these BRD4 Acyl-eCRs is specifically caused by the PROTAC-induced degradation, since treatment with JQ1, used by the ARV-825 PROTAC to target bromodomains, does not affect the protein abundance. Additionally, the ARV-825-induced degradation maintains specificity for its intended targets, as the mutant BRD4(2) BRD Acyl-eCR, is not affected by the PROTAC treatment, similar to the empty eGFP control (Figure 2b).

The JQ1 inhibitor, which is used as the warhead in the ARV-825 PROTAC, is known to have a preference for binding to the second bromodomain of BRD4 over the first bromodomain of BRD4^8^. This preference for the second bromodomain is also observed with the ARV-825 PROTAC, as the Acyl-eCR containing only the second bromodomain (BRD4(2).1x) is degraded more efficiently and faster than the construct containing the first bromodomain (BRD4(1).1x) (Figure 2c and Supplemental Figure 3a). In addition, we observe that increasing the valency of bromodomains in an Acyl-eCR construct increases the PROTACs ability to degrade it. This is seen by Acyl-eCRs with more copies of the second BRD4 bromodomain being degraded faster by ARV-825 (Figure 2c and Supplemental Figure 3a). Additionally, the BRD4(1_2).1x Acyl-eCR is more efficiently degraded by ARV-825 than constructs with only one copy of each bromodomain (Figure 2c and Supplemental Figure 3a). Together, these findings demonstrate that both bromodomain specificity and valency influence the efficiency of PROTAC-mediated degradation, and that Acyl-eCRs are suitable tools to measure these effects.

### Acyl-eCR-Based Competitive Binding Assays Distinguish Inhibitor Specificity Across Bromodomains

While Acyl-eCRs may not always be suitable for screening PROTAC specificity and potency, due to the lack of other protein domains that influence the binding and ternary complex formation with E3 ligase complexes^22^, they can still be employed for screening small molecule inhibitors that bind to the bromodomain. To take advantage of this, we created a treatment scheme where Acyl-eCRs are used to measure the binding of small molecule inhibitors *in cellulo* by competing with validated PROTACs for binding to the same bromodomain pocket (Figure 3a). We reasoned that if the small molecule inhibitor could strongly bind the bromodomain of interest, then any PROTAC-induced degradation of the Acyl-eCR would be reduced, thereby allowing for the measurement of the inhibitor’s binding strength.

**Figure 3.**
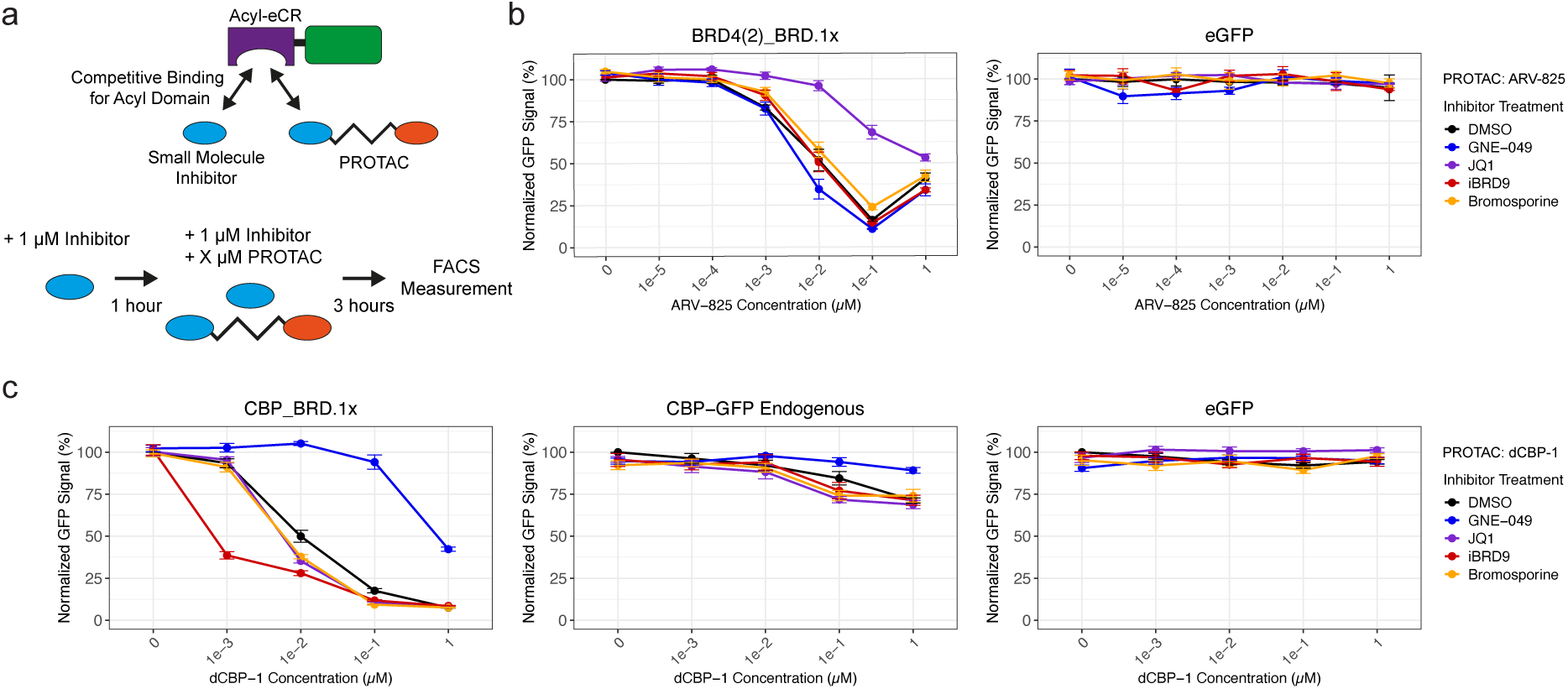
**(A)** *Top:* Schematic showing how the competitive binding of small molecule inhibitors versus PROTACs for the binding pocket of Acyl-eCRs can be used to measure the affinity of a small molecule for a bromodomain *in cellulo*. Inhibitors with higher affinity for a bromodomain, better prevent PROTAC-induced degradation. *Bottom*: Treatment scheme for competitive binding experiments. Cells were treated with 1 μM inhibitors for 1 hour. Then, varying concentrations of the PROTAC were added in addition to the previously added inhibitor. After 3 hours of treatment, the cell fluorescence was measured via flow cytometry. **(B)** Competitive binding between ARV-825 and several small molecule inhibitors showing how the inhibitors bind to BRD4(2)_BRD.1x. The cells were treated with the indicated inhibitor at a 1 μM concentration for 1 hour. Then, the stated concentration of ARV-825 PROTAC was added for 3 hours, in addition to the previous concentration of the same inhibitor. The percentage represents the GFP signal in treated cells as a ratio of the signal observed in untreated samples of the same cell type, after normalizing for the autofluorescence of the drug treatment in wild-type cells. **(C)** Competitive binding between dCBP-1 and several small molecule inhibitors showing how the inhibitors bind CBP bromodomains in Acyl-eCR constructs versus the endogenous CBP protein. The cells were treated with the indicated inhibitor at a 1 μM concentration for 1 hour. Then, the stated concentration of dCBP-1 PROTAC was added for 3 hours, in addition to the previous concentration of the same inhibitor. The percentage represents the GFP signal in treated cells as a ratio of the signal observed in untreated samples of the same cell type, after normalizing for the autofluorescence of the drug treatment in wild-type cells.

To test this competitive binding treatment scheme, the BRD4(2).1x Acyl-eCR cell line was first treated for one hour with several small molecule inhibitors specific to BRD4, CBP/p300, BRD9, and a broad-spectrum bromodomain inhibitor (Bromosporine) at 1 µM concentration to allow engagement with the bromodomain. After 1 hour, the BRD4 specific PROTAC, ARV-825, was added at increasing concentrations for 3 hours in the presence of the small molecule inhibitors, followed by measurements via flow cytometry. This showed that the CBP/p300 inhibitor GNE-049, the BRD9 specific inhibitor iBRD9, and the broad-spectrum bromodomain inhibitor Bromosporine, did not block the ARV-825 induced protein degradation, resulting in >50% GFP reduction already in the presence of 10nM of the PROTAC compound (Figure 3b). However, the BRD4 specific inhibitor, JQ1, did block the PROTAC-induced degradation and required equimolar concentrations of ARV-825 to reduce GFP levels to 50% (Figure 3b). The same competitive binding assay was repeated using the CBP.1x Acyl-eCR, where GNE-049, a CBP/p300-specific inhibitor, blocked dCBP-1-mediated degradation with similar kinetics (Figure 3c). Similar results were observed for the endogenously GFP-tagged CBP protein, but to a lesser extent, since the dCBP-1 PROTAC only partially degrades the full-length protein in these conditions (Figure 3c).

To further interrogate inhibitor binding, we reversed the treatment scheme, keeping the PROTAC concentration constant while varying the inhibitor concentration (Supplementary Figure 4a). This showed that an equimolar dose of GNE-049 effectively reduced dCBP-1– induced degradation of the CBP.1x Acyl-eCR by 50%, while the other inhibitors had no effect (Supplementary Figure 4b). This modified scheme provides additional resolution by quantitatively estimating inhibitor affinity within cells, which is not captured when only PROTAC levels are titrated. Together, these two complementary treatment schemes allow both qualitative validation of inhibitor specificity and quantitative estimation of bromodomain binding affinity, using Acyl-eCRs as modular reporter constructs.

### A panel of Acyl-eCR expressing cell lines allows comparative and quantitative analysis of small molecule specificity towards bromodomains from various proteins

To evaluate the broader applicability of this competitive binding assay, additional Acyl-eCR cell lines were generated, comprising a total of 14 different bromodomains. This panel of cell lines represents 14 different proteins and was selected to represent the different types of structural and druggable characteristics of bromodomains^23^ (Figure 4a). All Acyl-eCR cell lines stably expressed the GFP fusion protein at high levels, maintaining consistent expression over several weeks in culture (Supplementary Figure 5a). To determine whether this panel can be used to test PROTAC-mediated degradation of bromodomains from non-target proteins, dCBP-1 PROTAC treatments were applied to all Acyl-eCRs cell lines, followed by GFP fluorescence measurements. While we observe efficient degradation of the CBP and p300 Acyl-eCRs, this assay shows that dCBP-1 is capable of partially degrading Acyl-eCRs from non-target proteins at varying efficiency (Figure 4b). Among the degraded Acyl-eCR, we observe BRD9, BRD7, and CECR2 showing >25% degradation (Figure 4b). Structural analysis of the bromodomains revealed that most constructs share moderate to high structural similarity with CBP or p300 (TM-score 0.78–0.9 and RMSD values < 2 Å), indicating that the panel of Acyl-eCRs provides a broad coverage of the bromodomain family tree (Supplementary Figure 5b and c). Comparing the structural similarity of each bromodomain in the Acyl-eCRs with the bromodomains of CBP or p300 revealed no clear correlation with the degradation efficiency, indicating that structural similarity alone does not predict susceptibility to dCBP-1–mediated degradation (Supplementary Figure 5b and c). These results demonstrate how this panel of Acyl-eCRs can be used to determine the *in cellulo* specificity & potency of available or novel PROTACs, extending the applicability of Acyl-eCRs.

**Figure 4.**
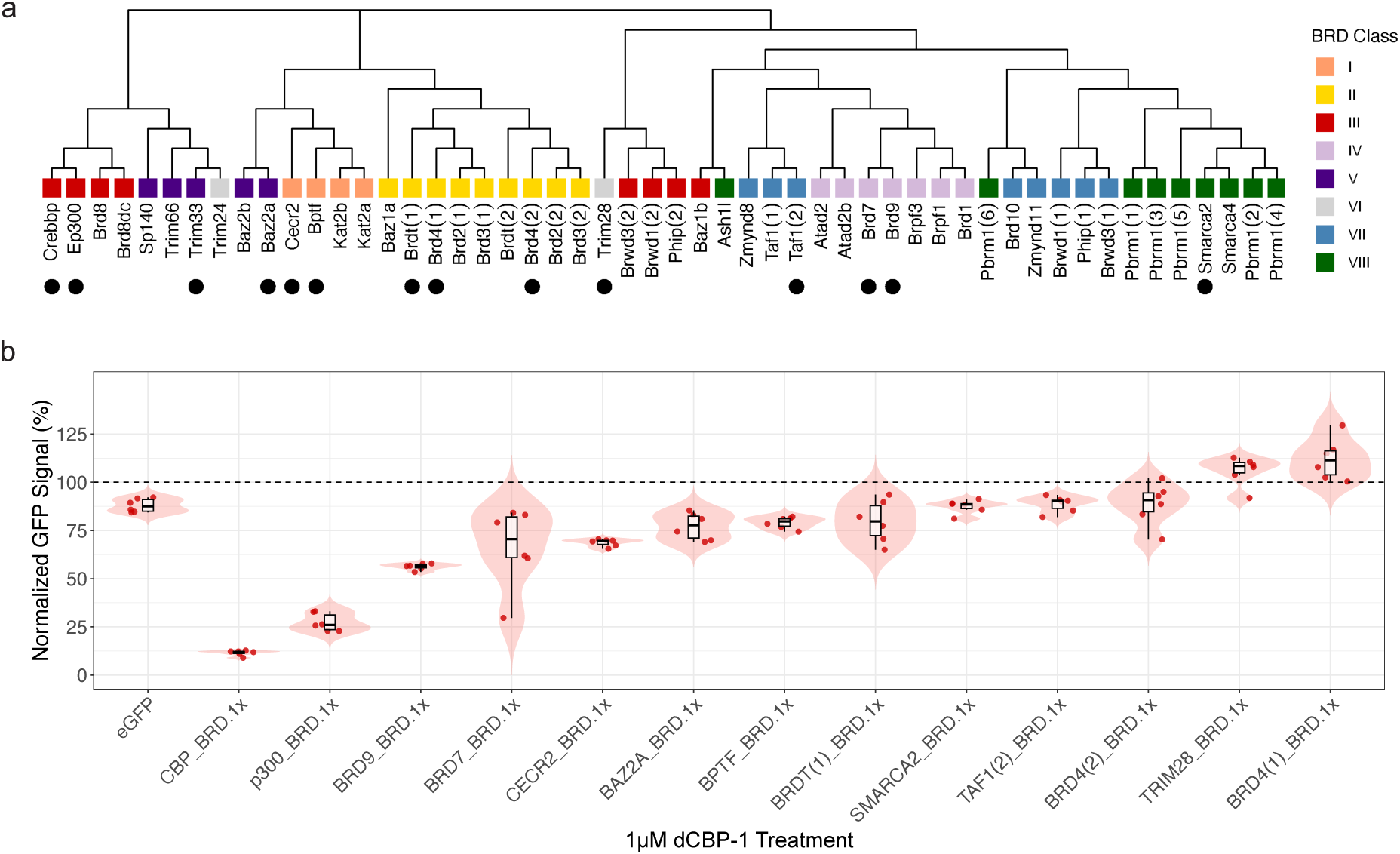
**(A)** Phylogenetic tree showing that the panel of Acyl-eCRs comprises representative bromodomains from all classes of bromodomains. All bromodomain protein sequences were obtained from InterPro^28^ and aligned with Clustal Omega’s^30^ multiple sequence alignment. Bromodomains are classified based on structure & druggability^23^. Black circles represent the bromodomains that are included in the panel of Acyl-eCR cell lines. **(B)** Normalized FACS data showing the effects of dCBP-1 PROTAC treatments on all Acyl-eCR cell lines. PROTAC was added at a 1 μM concentration for 24 hours of treatment. The percentage represents the GFP signal in treated cells as a ratio of the signal observed in untreated samples of the same cell type, after normalizing for the autofluorescence of the drug treatment in wild-type cells.

### Nuclear retention of Acyl-eCRs measures small-inhibitor effectiveness in cells

To establish a method for screening small-molecule inhibitors independent of PROTAC-based degradation of Acyl-eCRs, we tested if the chromatin retention of Acyl-eCRs after inhibitor treatment could be used to assess target engagement *in cellulo*. To do this, we adapted and extended existing protocols designed to quantify the nuclear binding of proteins of interest using flow cytometry^24,25^. Here, fluorescence measurements taken after the extraction and permeabilization of nuclei allow for the quantification of the retained chromatin-bound fraction of Acyl-eCRs, while the unbound fraction is washed out (Figure 5a). By measuring the level of signal retained in the nucleus following drug treatment and protein wash-out, we can infer the extent to which a compound disrupts the interaction between the Acyl-eCR construct and its nuclear binding target.

**Figure 5.**
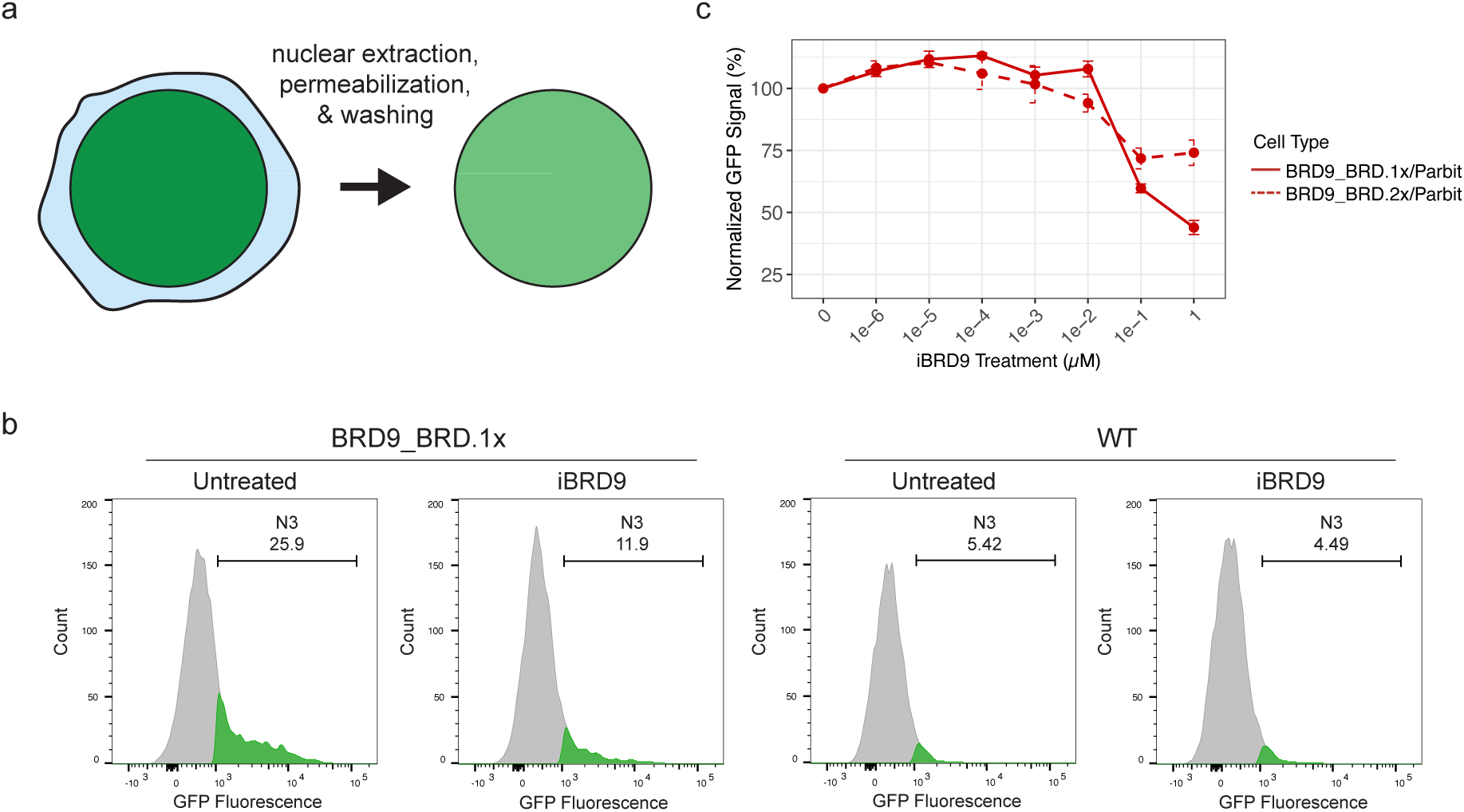
**(A)** Schematic showing how nuclei isolation & permeabilization followed by flow cytometry can measure the retention of proteins on chromatin in lieu of high background fluorescence. Nuclei can be harvested and permeabilized from whole cells, and then washed to remove the unbound or weakly bound fraction of the protein of interest. Since the weakly bound fraction of protein is removed, the fraction of protein remaining can be measured at a better signal-to-noise ratio via flow cytometry. **(B)** Flow cytometry analysis of nuclei harvested from BRD9_BRD.1x or WT cells. N3 gate shows the GFP signal being measured in nuclei, after iBRD9 or control treatments. Treatments in the WT cell line show a change in autofluorescence in the nuclei from the drug treatments. **(C)** Normalized flow cytometry data showing how Acyl-eCRs with 1x or 2x copies of the BRD9 bromodomain remain bound to chromatin, after iBRD9 treatments. iBRD9 treatments were performed at 1 μM concentration for 24 hours. The percentage represents the GFP signal in treated cells as a ratio of the signal observed in untreated samples of the same cell type, after normalizing for the autofluorescence of the drug treatment in wild-type cells.

To test this approach, we selected cells expressing the BRD9_BRD.1x Acyl-eCR and treated them with iBRD9^26^ at a concentration of 1 μM for 24 hours. Nuclei were harvested and permeabilized with a mild extraction buffer, and fluorescent signals from the nuclear-retained Acyl-eCRs were obtained by gating for single-nuclei populations (Supplemental Figure 6a). Nuclear retention of the BRD9_BRD.1x Acyl-eCR without any inhibitor treatments showed a fraction of cells with a higher fluorescent signal, where the Acyl-eCR maintained binding in the nucleus (Figure 5b). This untreated condition of cells was used as a baseline, and an arbitrarily drawn gate, N3, was drawn to represent the cells in which a high signal is maintained. Upon treatment with iBRD9, the proportion of cells within the N3 gate decreased from 25.9% to 11.9% of the total population, indicating effective displacement of the Acyl-eCR construct from chromatin and the nucleus (Figure 5b). Replicates from this assay also gave similar results, indicating that the measurements are robust despite the nuclear permeabilization & washing steps (Supplementary Figure 6b).

By repeating this nuclear retention assay with Acyl-eCRs containing multiple copies of the BRD9 bromodomain and with increasing concentrations of iBRD9, we found that the Acyl-eCR with two bromodomains was retained more effectively in the nucleus at 1 μM concentrations than the Acyl-eCR containing only one domain. This demonstrates that increasing the number of bromodomain copies enhances chromatin binding stability and may buffer against displacement by small-molecule inhibitors (Figure 5c). Together, these findings highlight the utility of the nuclear-retention cytometry assay as a quantitative approach for assessing inhibitor potency and target affinity in a cellular context.

## Discussion

Here, we present a panel of cell lines expressing individual and combinations of bromodomains as a versatile platform to measure the potency and specificity of small-molecule inhibitors in living cells. By implementing two complementary flow cytometry–based assays, we demonstrate that these synthetic constructs can report on inhibitor-target engagement without requiring the full-length endogenous protein. This approach helps address a major bottleneck in chemical biology: the frequent disconnect between *in vitro* measurements of compound activity and their actual behavior in cells. Many drug candidates that show promising binding affinities *in vitro* fail to demonstrate efficacy in *vivo* due to issues such as poor permeability, lack of target accessibility, unexpected off-target interactions, post-translational modifications, or other context-dependent reasons. Our cellular assays help address this challenge by offering scalable, direct readouts of inhibitor-target engagement under physiological conditions.

In the first approach, we used competitive binding assays to assess the affinity and specificity of small-molecule inhibitors by leveraging the off-target degradation capacity of certain PROTACs. Although PROTACs are typically designed to degrade specific proteins, many exhibit broader degradation profiles due to binding promiscuity and the influence of additional structural features from other protein domains that facilitate ternary complex formation. We took advantage of this property to use PROTAC-induced degradation of Acyl-eCRs as a readout for inhibitor competition: if an inhibitor binds to the same bromodomain as a PROTAC, it can protect the Acyl-eCR from degradation. Using this logic, we could quantify inhibitor affinity by titrating either the PROTAC or the inhibitor concentration. However, this strategy is inherently dependent on whether a given PROTAC can degrade the bromodomain construct in question. Since PROTAC degradation often requires additional context from the full-length protein, such as neighboring domains or specific surface interactions, not all bromodomain constructs are susceptible to off-target degradation. As a result, the competitive binding assay is only applicable to a subset of Acyl-eCRs where degradation by the given PROTAC is sufficiently robust and specific. This limits its applicability for drug screening across a broader panel of domains.

To overcome this limitation, we developed a second approach based on nuclear-retention FACS assays, which measure chromatin association of bromodomain constructs in response to inhibitor treatment. In this method, nuclei are permeabilized to wash out unbound proteins, and the remaining nuclear-retained GFP signal is quantified by flow cytometry. A decrease in fluorescence upon drug treatment reflects displacement of the bromodomain from the nucleus, offering a direct, degradation-independent readout of target engagement. This assay builds upon concepts previously used in FRAP (fluorescence recovery after photobleaching) studies, which have measured chromatin residence time of bromodomain proteins as a proxy for inhibitor binding. FRAP has been the standard method for evaluating bromodomain inhibitor activity in cells, and more broadly for assessing small-molecule engagement of many chromatin reader domains^21^. However, unlike FRAP, which is low-throughput and requires specialized microscopy, this nuclear-retention FACS approach is rapid, scalable, and amenable to multi-well screening formats. That said, it may not be suitable for all bromodomains, as domains with low-affinity or transient chromatin interactions may not retain sufficient signal for quantification.

Both approaches presented here are valuable tools for the *in cellulo* analysis of bromodomain inhibitors. Notably, neither method requires recombinant protein purification or in vitro biochemical characterization, making them especially attractive for early-stage drug screening or compound triaging. Additionally, they can be applied even when the biological function or chromatin targets of the bromodomain are unknown, making them useful for understudied or non-canonical reader proteins. Beyond bromodomains, the Acyl-eCR strategy and these FACS-based assays could be extended to other epigenetic reader domains, such as chromodomains, PHD fingers, or Tudor domains. As long as the fusion construct shows measurable nuclear binding and responds to small-molecule perturbation, the system can provide a live-cell proxy for binding affinity and specificity. In this way, our work establishes a generalizable platform for the functional evaluation of chromatin-targeting drugs in a native cellular context.

## Materials & Methods

### Cell culture and drug treatments

Mouse embryonic stem cells (HA36CB1) were cultured on 0.2% gelatin-coated plates. Cells were grown in media containing DMEM (Invitrogen) that was supplemented with 15% fetal calf serum (Invitrogen), 1x non-essential amino acids (Invitrogen), 1x glutamax (Invitrogen), 0.001% 2-mercaptoethanol (Invitrogen), and leukemia inhibitory factor (LIF) (made in-house) at 37°C in 5% CO_2_.

The CBP/p300 bromodomain inhibitor: GNE-049 (MedChemExpress, HY-108435), CBP/p300 PROTAC: dCBP-1 (MedChemExpress, HY-134582), BRD4 bromodomain inhibitor: (+)-JQ-1 (MedChemExpress, HY-13030), BRD4 PROTAC: ARV-825 (MedChemExpress, HY-16954), BRD9 bromodomain inhibitor: iBRD9 (MedChemExpress, HY-18975), and broad-spectrum bromodomain inhibitor: Bromosporine (MedChemExpress, HY-15815) were dissolved in DMSO and then diluted to 1μM in mESC media for 24-hour treatments, unless stated otherwise.

### Cell line generation

Acyl-eCR cell lines were generated via RMCE, as previously described^13^. In brief, bromodomains were amplified from cDNA or synthesized (Integrated DNA Technologies) based on domain annotations (Uniprot). Bromodomain coding sequences were cloned into the parbit-v6 plasmid (Addgene #179392) via Gibson assembly. The resulting construct has a constitutively active CAG promoter that drives the expression of the bromodomain of interest fused to a nuclear localization signal, eGFP, an internal ribosome entry site, and a puromycin-*N*-acetyltransferase resistance cassette. Each of these RMCE constructs were co-transfected with a CRE recombinase expression plasmid (Addgene #19131) in a 1:0.6 DNA ratio using Lipofectamine 3000 (L3000015, Thermo Fisher Scientific). A double-selection was performed to generate a homogeneous population of cells expressing each Acyl-eCR of interest. First, 3μM Ganciclovir (Selleckchem, #S1878) was applied for 4 days, followed by 2μM Puromycin (Gibco, A1113803) for 2 days. Flow cytometry was performed to check for the stable expression of the eGFP fusion and to ensure that the cells were homogeneously expressing the construct of interest.

Cell lines where CBP or p300 are endogenously tagged with eGFP were generated by co-transfecting sgRNAs targeting the C-terminus of either CBP or p300 in the pX330-U6-Chimeric_BB-CBh-hSpCas9 backbone (Addgene #42230) (Table 2), pRR-Puromycin recombination reporter plasmids (Addgene #65853) for each sgRNA (Table 2), and a donor plasmid for homology-directed repair (Table 2). These donor plasmids consisted of an eGFP coding sequence followed by a stop codon flanked by sequences homologous to the C-terminus of either CBP or p300. After co-transfection, 2μM Puromycin (Gibco, A1113803) was added for 36 hours. Clones positive for the homologous eGFP knock-in were picked, genotyped, and validated via Sanger sequencing and western blotting.

### Western blotting

For Acyl-eCR and endogenously tagged-eGFP detection, crude nuclear extracts were prepared from cells as previously described^27^, and 20 μg were run on 4-12% NuPAGE gels (Thermo Fisher Scientific). Proteins were transferred to PVDF membranes and then blocked with 5% (wt/vol) milk in TBS-0.1% Tween-20. Membranes were incubated overnight at 4°C with primary antibodies for GFP (Abcam, ab290). Protein detection was done using species-specific antibodies conjugated to horseradish peroxidase and Pierce® Peroxidase IHC Detection Kit (Thermo Scientific). Total protein loading was assessed using Revert™ 700 Total Protein Stain (LI-COR Biosciences, 926-11011) according to the manufacturer’s instructions.

### Flow cytometry measurements and analysis

Cells were seeded onto 96-well plates (TPP, 92196) prior to treatments and measurements. Cells were seeded such that there would be 20,000 cells in each well on the day of measurement. After treatments, cells were washed with PBS (Gibco, 10010023) and trypsinized (Gibco, 15400054). The trypsin was then blocked with DMEM with 10% FCS. Cells were then analyzed on a FACSymphony A1 Analyzer with a High Throughput Sampler (BD Biosciences). Cells were gated for individual and viable cells, as well as channel voltages for GFP (Alexa Fluor 488-A), with respect to the validated negative and positive eGFP expressing controls. Analysis and visualization were performed using FlowJo (Version 10.0.7, TreeStar). The geometric mean of the eGFP signal was exported and used for analysis in custom scripts.

A two-step normalization approach was employed to account for variations in eGFP signal strength across different cell lines and to correct for the autofluorescence associated with the different drug treatments. First, the geometric mean fluorescence intensity of the eGFP signals from each cell line of interest was adjusted relative to the WT cells of the same drug treatment, to normalize for inherent differences in expression of each eGFP fused bromodomain. Specifically, the eGFP signal from each treated condition of the eCR cells was normalized by subtracting the eGFP signal from the untreated WT cells; effectively controlling for baseline autofluorescence. Then, a ratio-based normalization was performed to quantify the effect of each treatment relative to a DMSO control treatment. This was calculated using the following formula:

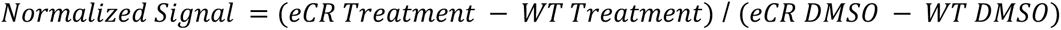

### Competitive binding assays

Competitive binding between PROTACs and a small molecular inhibitor for the binding pocket of bromodomains was done via a two-part treatment approach. First, the small molecule inhibitor was added to the cells at the stated concentration for 1 hour. After this time had elapsed, all of the media was removed from the cells and replaced with media containing the stated concentrations of the small molecule inhibitor and the PROTAC. This dual treatment was performed for 3 hours. Afterwards, the cells were harvested for flow cytometry analysis.

### Immunofluorescence microscopy

mESCs were cultured on sterile coverslips (0.13–0.16 mm – Ted Pella) pre-coated with 0.2% gelatin. After treatment, cells were fixed with 2% methanol-free formaldehyde (Thermo Fisher Scientific, 28908) in PBS for 15 min at RT. Cells were washed three times with PBS for 5 min at RT and stained with Hoechst (Thermo Fisher Scientific, 62249) for 10 min. Coverslips were then washed three times for 5 min with 1X PBS and mounted on glass slides with 5 μl Mowiol-based mounting media (Mowiol 4.88, Calbiochem, #475904) in Glycerol/TRIS. Images were acquired with a Zeiss LSM 700 confocal laser scanning microscopy (Biology Image Center, Utrecht University, Netherlands) with a Plan-Apochromat 63X 1.4 NA oil immersion objective (Carl Zeiss, Germany) using the 405 and 488-nm lasers. Images consist of a z-stack of 5-8 planes at 0.37 µm intervals. Maximum Intensity Z-projections of Z-stacks were implemented with Fiji (version2.14.0/1.54f). The bottom panel of Figure 2b was also implemented using Fiji. A panel of representative eGFP fluorescence images, acquired with the same microscope settings, was pseudocolored using the 16-color lookup table (LUT), and a calibration bar indicating pixel counts for each gray value in the 16-bit images was then generated. The mean intensity of GFP signal was determined with CellProfiler (Broad Institute, version 4.2.6) (Stirling et al., 2021). Images were corrected for illumination biases using the “Rescale Intensity” module. A Gaussian filter was applied to remove the foreground signal. Nuclei (Hoechst stained) were segmented with the Otsu threshold and watershed modules to produce a binary image. Segmented nuclei were further identified using the “IdentifyPrimaryobjects” module. Missegmented nuclei were manually inspected and further discarded from the analyses. The object intensity (488nm) was then obtained for each segmented nucleus.

### Phylogenetic tree

All mouse bromodomain proteins were obtained from InterPro^28^ using the Pfam^29^ ID #PF00439. Protein sequences were downloaded, and only the InterPro-reviewed sequences were kept. The Pfam annotation was then used to obtain the protein sequence of each bromodomain. If a given protein had multiple bromodomains, each instance’s protein sequence was obtained and numbered.

The protein sequences of each mouse bromodomain underwent a multiple sequence alignment (MSA) using Clustal Omega^30^. The resulting tree was then plotted in R using the “ggTree”^31^ package. The bromodomains were then colored based on their structure & drugability scores^23^.

### Structural comparisons

The amino acid sequences for the 14 bromodomains incorporated into the Acyl-eCR panel were identified based on their corresponding Pfam annotations^29^. To characterize the structural features of these bromodomains, their three-dimensional structures were predicted using ColabFold^32^. The highest-ranked predicted structure for each bromodomain was selected for subsequent analysis.

To provide a common reference for structural comparison, all predicted bromodomain structures were aligned to the CBP or p300 bromodomains, which were used as reference structures. This alignment was performed using the TM-align (Template Modeling alignment) algorithm^33^. The quality of the structural alignments was quantified by calculating both TM-scores and Root-Mean-Square Deviation (RMSD) scores for each bromodomain relative to the reference bromodomain, using US-align^34^.

For each bromodomain, degradation efficiency was determined from FACS-based measurements of the normalized GFP signal. Replicate measurements were averaged, and these mean values were used for subsequent correlation analyses. Pearson correlation coefficients (r) were calculated between degradation efficiency and either RMSD or TM-scores, relative to the CBP or p300 bromodomains. To visualize these relationships, scatterplots were generated showing the mean ± SD for each bromodomain, along with a linear regression line and annotated correlation coefficients.

### Nuclear retention FACS assay

The quantification of nuclear retained Acyl-eCR proteins was performed using a modified flow cytometry-based chromatin retention assay, adapted from previously described protocols^24,25^. Cells were seeded and treated with small-molecule inhibitors as indicated. 2.5x10^5^ cells were then washed with PBS to remove debris, trypsinized to create a single-cell suspension, and the reaction was stopped by adding TrypStop (DMEM (Invitrogen) with 10% FCS (Invitrogen)) to neutralize the trypsin. The cells were then centrifuged at 200×g for 5 minutes, the supernatant was aspirated, and the cell pellet was resuspended in 100 µL CSK buffer (25 mM HEPES pH 7.4, 50 mM NaCl, 1 mM EDTA, 3 mM MgCl2, 300 mM sucrose, 0.5% (v/v) Triton X-100, and a complete protease inhibitor cocktail (Roche, 11697498001)) for 10 minutes on ice to lyse cells and permeabilize nuclei. Following lysis, nuclei were washed three times with PBS-B (1 mg/ml BSA in 1× PBS) by centrifuging the cells at 400×g for 5 minutes and removing the supernatant. Nuclei were then resuspended in analysis buffer (0.02% (w/v) sodium azide in PBS-B) for storage and analysis.

Samples were analyzed by flow cytometry. Single-cell populations were identified by gating on FSC-A versus SSA-A plots. The nuclear-retained GFP fluorescence signal was then measured for each sample. For each cell line, an arbitrary gate was established on the untreated control sample to define the high-fluorescence, chromatin-bound population. The percentage of cells falling within this gate was used to quantify the extent of chromatin retention, and this value was compared across different inhibitor treatments to assess their effectiveness at displacing Acyl-eCRs from chromatin.

## Acknowledgements

We thank members of the Baubec laboratory for their input and criticism. Furthermore, we thank Toni van Capel from the Utrecht University FACS facility and members of the Biology Imaging Center at Utrecht University for their support. This work was supported by the European Research Council (865094 - ChromatinLEGO - ERC-2019-COG) and by the Swiss National Science Foundation through SNSF Sinergia (180354).

## Author contributions

D.C.R., with the help from R.C.S and N.K., performed all experiments and generated reagents. A.T., S.G., and R.V. generated constructs and cell lines. D.C.R. and T.B. analyzed the data and wrote the manuscript with input from all authors.

## Supplemental Figures and Legends

**Figure S1.**
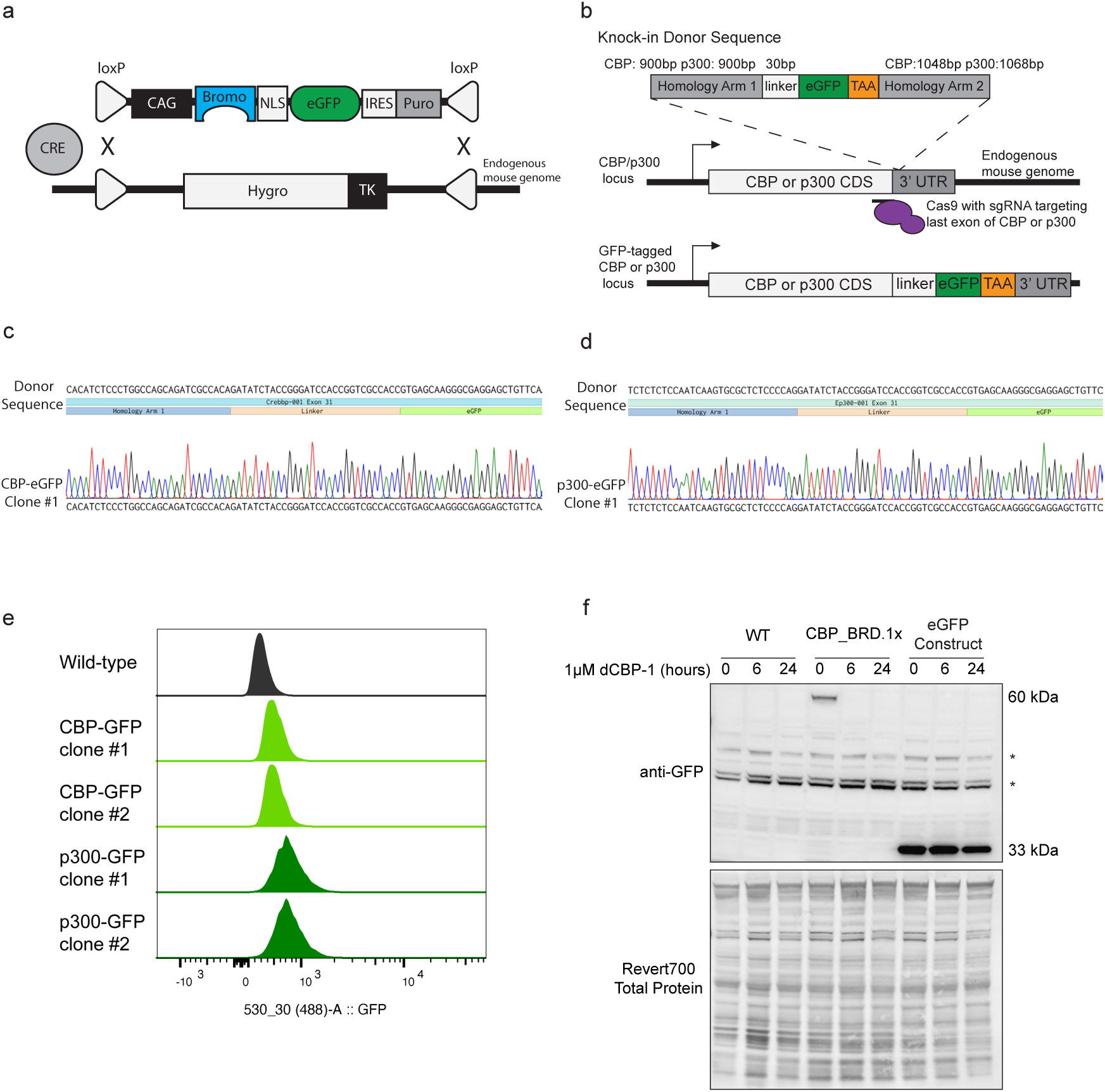
**(A)** Schematic showing how the Parbit expression cassette is used to generate stably expressed Acyl-eCRs in mESCs. RMCE by Cre, followed by a double selection of ganciclovir and puromycin, was applied to generate these constructs at a defined site in the mouse genome. A CAG promoter drives the constitutive expression of the bromodomain of interest, which is fused to a nuclear localization signal (NLS), and an eGFP tag. This construct also fuses a biotin acceptor site to the N-terminus of the protein, which can be biotinylated in vivo by a bacterial BirA ligase. **(B)** Schematic diagram showing how CBP was endogenously tagged with an eGFP tag. A homology donor construct was generated by cloning a 900 bp upstream and a 1,048 bp (CBP) or 1,068 bp (p300) downstream homology arm flanking a 30 bp flexible GGS linker that was fused to an eGFP tag. This donor construct was co-transfected with a pX330 CRISPR-Cas9 plasmid, which had an sgRNA targeting the C-terminus of the CBP gene. **(C-D)** Sanger sequencing of genotyping PCR products from the C-terminus of the CBP **(C)** and p300 **(D)** loci, confirming the in-frame homologous integration of an eGFP tag. Data shown are from mESC clone #1 for both CBP and p300 tagging. **(E)** Flow cytometry data showing the eGFP signal from cell lines where either CBP or p300 were endogenously tagged with eGFP. Two clonal replicates for each protein tagging are shown. Cell lines were maintained in culture for more than two weeks to demonstrate stable expression of the eGFP fusion proteins. **(F)** Western blot of nuclear extracts from cell lines treated with 1 μM dCBP-1 PROTAC for the stated duration. An antibody against GFP was used to probe the eGFP tag on the Acyl-eCR constructs. The CBP_BRD.1x eCR runs at approximately 60 kDa, and the Empty-eGFP construct runs at 33 kDa. Non-specific bands are marked by an asterisk (*). Revert700 Total Protein Stain is used to show equal loading in lanes.

**Figure S2.**
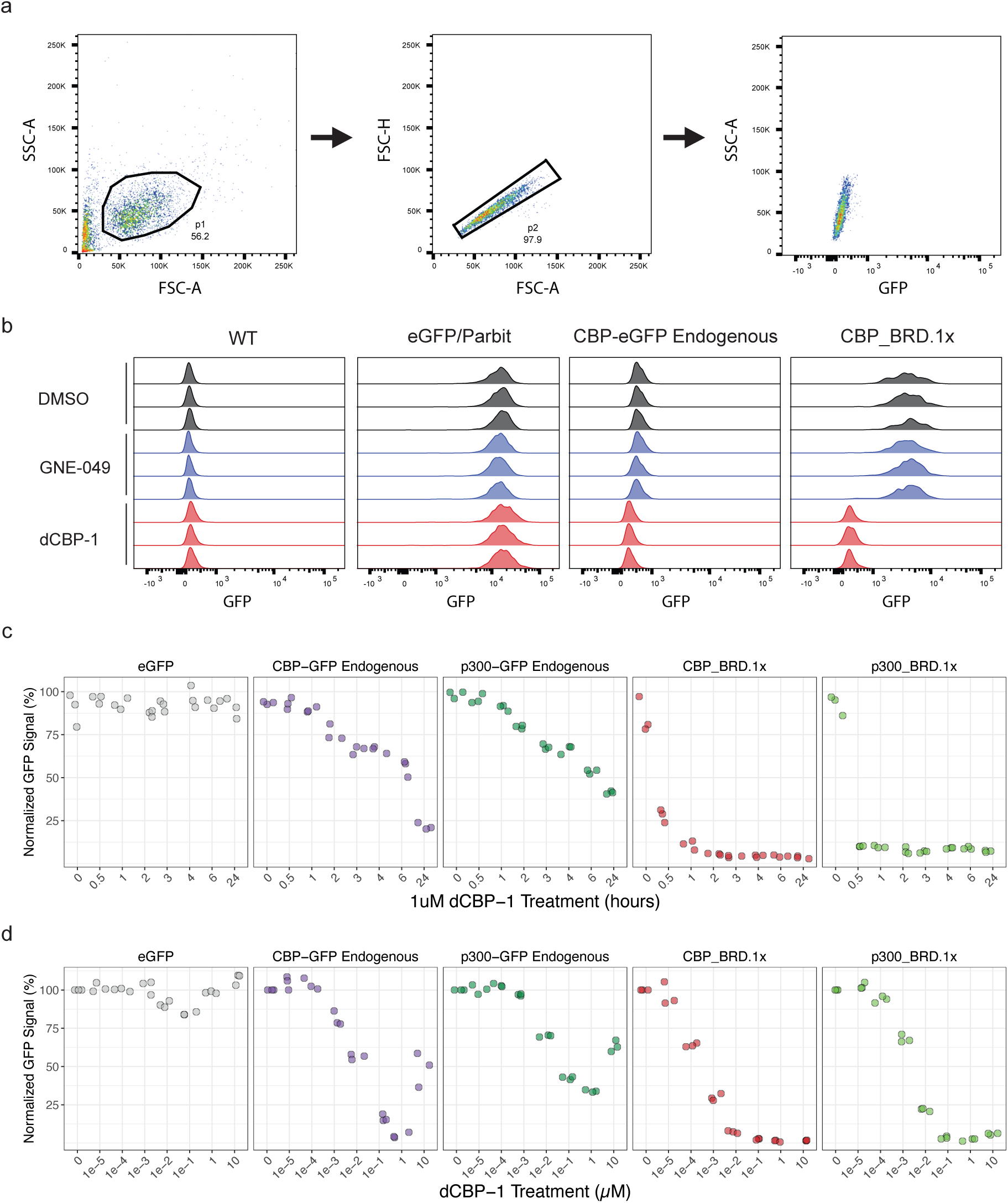
**(A)** Gating strategy for whole cell flow cytometry. Whole mESC populations were initially gated in region p1. Single cells were subsequently gated in region p2 to exclude doublets and debris. eGFP fluorescence signal was collected using the 530/30 nm bandpass filter with 488 nm excitation. **(B)** Flow cytometry analysis of eGFP fluorescence in multiple cell lines after different drug treatments. Three randomly selected replicates per treatment condition are shown for each cell line to show variability amongst replicates **(C)** Flow cytometry analysis of dCBP-1 PROTAC treatment over multiple timepoints at a fixed concentration of 1 μM. GFP signal is reported as a percentage relative to untreated cells of the same type, after normalization for drug-induced autofluorescence in wild-type cells. **(D)** Flow cytometry analysis of dCBP-1 PROTAC treatment at increasing concentrations, following 24-hour exposure. GFP signal is shown as a percentage of the signal in untreated cells of the same type, normalized to account for autofluorescence from the drug treatment in wild-type controls.

**Figure S3.**
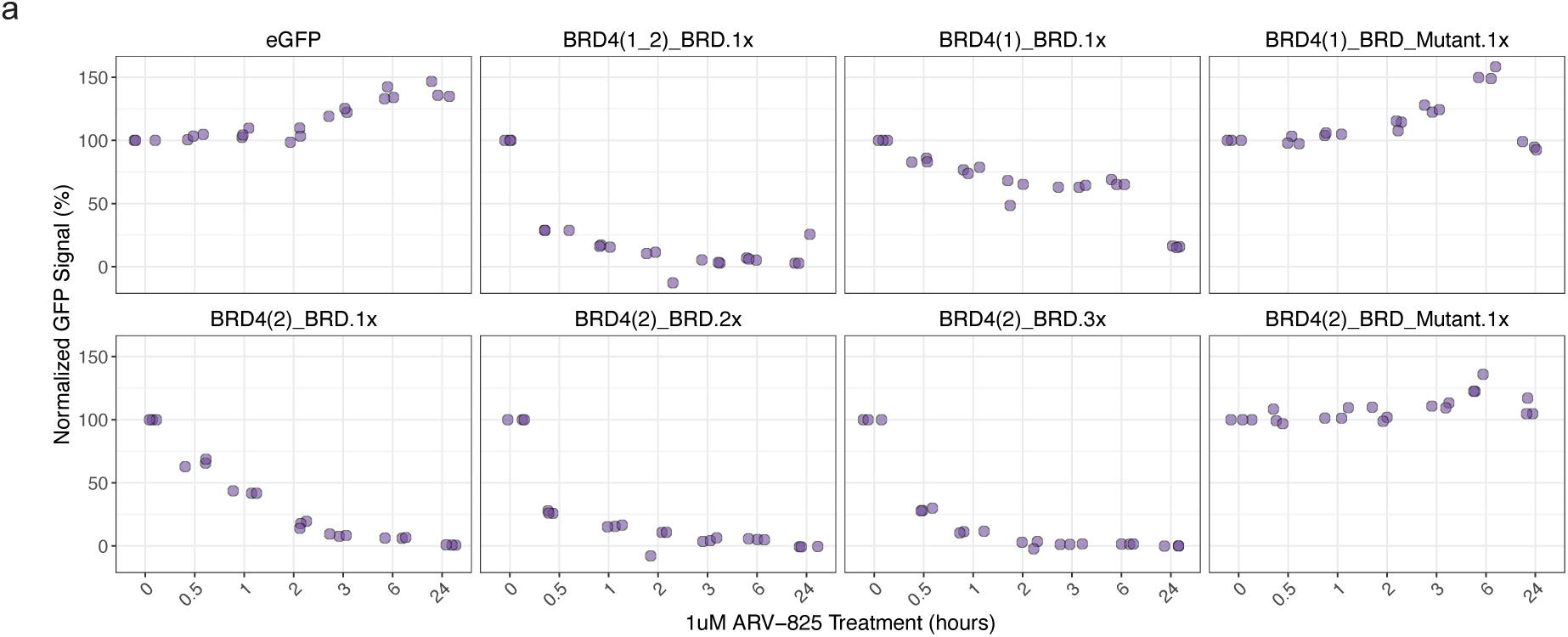
**(A)** Normalized FACS data showing the effects of ARV-825 PROTAC treatment over multiple time points on cells expressing various combinations of bromodomains from BRD4. All ARV-825 treatments were performed at a concentration of 1 μM. The percentage represents the GFP signal in treated cells as a ratio of the signal observed in untreated samples of the same cell type, after normalizing for the autofluorescence of the drug treatment in wild-type cells.

**Figure S4.**
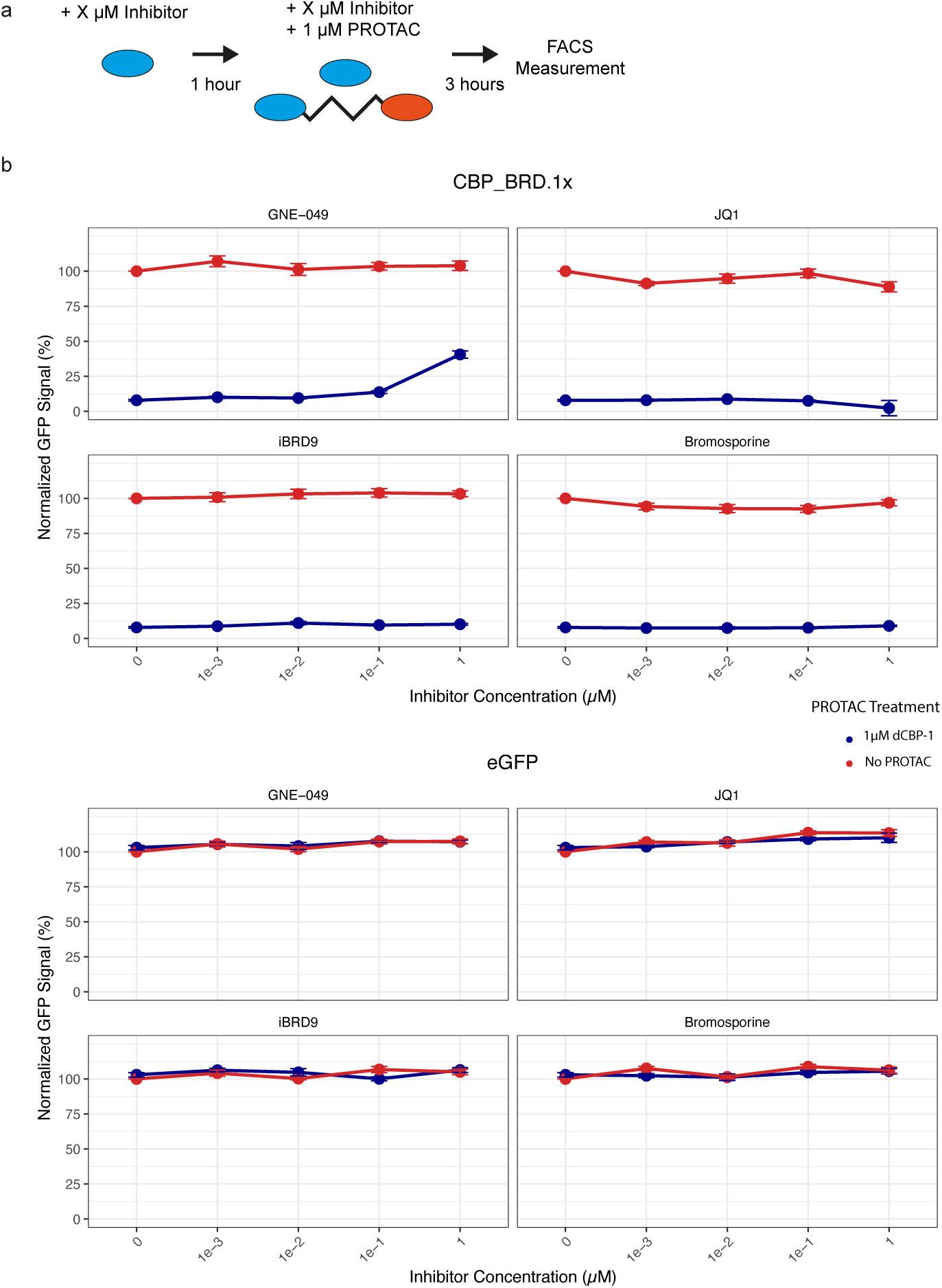
**(A)** Diagram showing the treatment scheme for this competitive binding experiment. Cells were treated with inhibitors for 1 hour at a concentration range from 1 μM to 0.001μM. Then, 1 μM of the dCBP-1 PROTAC was added in addition to the previously added inhibitor. After 3 hours of treatment, the cell fluorescence was measured via flow cytometry. **(B)** Competitive binding between dCBP-1 and several small molecule inhibitors showing how the inhibitors bind CBP bromodomains in Acyl-eCR constructs versus the endogenous CBP protein. The cells were treated with the indicated inhibitors for 1 hour, at the stated concentrations. Then, 1 μM of dCBP-1 PROTAC was added for a 3 hour treatment, in addition to the previous concentration of the same inhibitor. The percentage represents the GFP signal in treated cells as a ratio of the signal observed in untreated samples of the same cell type, after normalizing for the autofluorescence of the drug treatment in wild-type cells.

**Figure S5.**
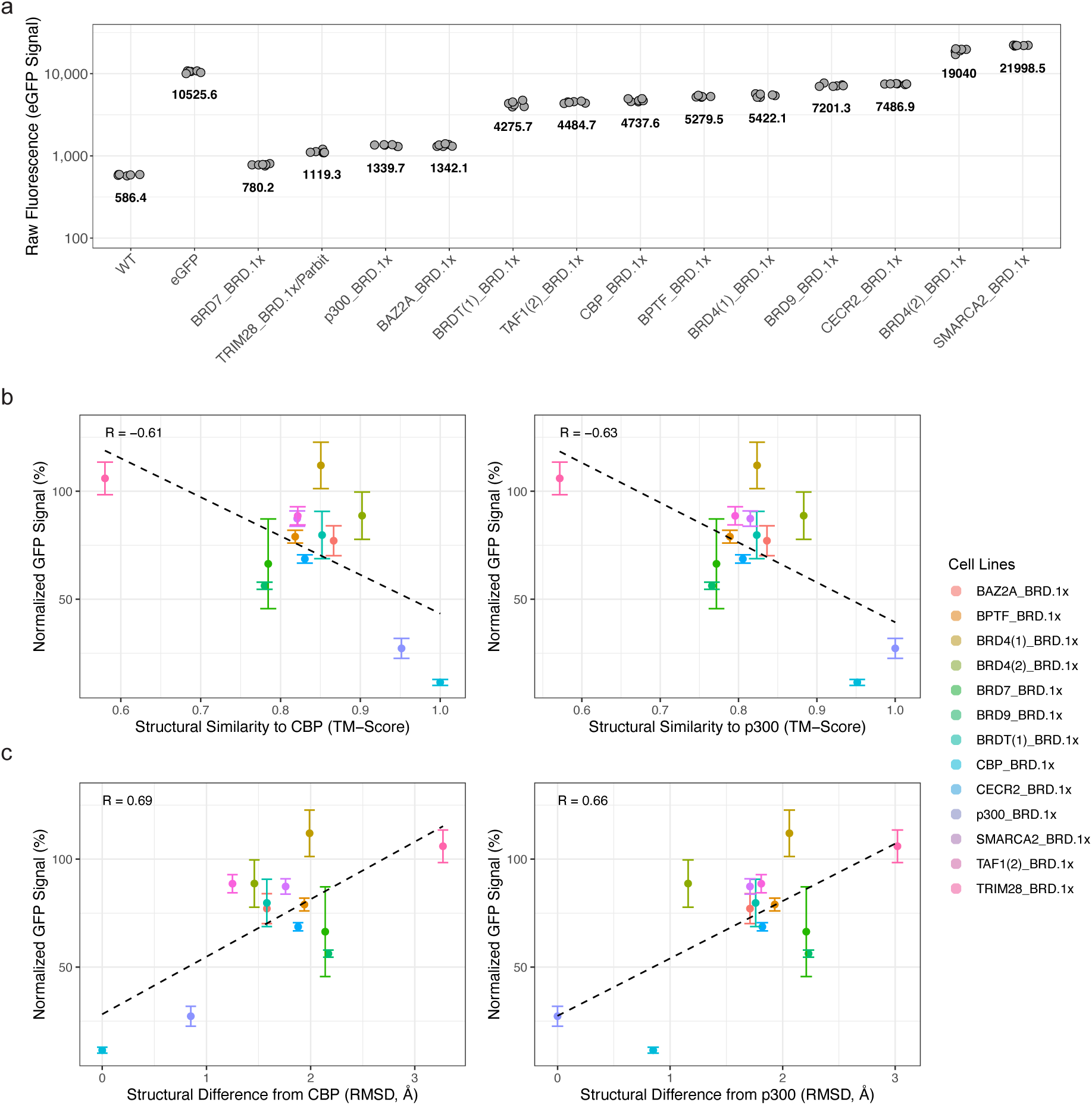
**(A)** Raw eGFP fluorescence intensity was measured in untreated cell lines expressing different acyl-eCR constructs after extended passaging to assess stable expression. Each dot represents an independent replicate measurement (n = 6 per construct). Black numbers denote the mean fluorescence intensity for each cell line. The y-axis is shown on a logarithmic scale. **(B-C)** Correlation between dCBP-1–mediated degradation of Acyl-eCR constructs and the structural similarity of their bromodomains to CBP or p300. Degradation efficiency (y-axis) was measured by flow cytometry as normalized GFP signal (%) after dCBP-1 treatment (data from Figure 4A; n = 6 independent measurements per construct). Structural similarity to CBP/p300 bromodomains was quantified using **(B)** TM-score and **(C)** RMSD (root-mean-square deviation). Each data point represents the mean ± SD of six replicates for a single construct, with a linear regression line and corresponding Pearson correlation coefficient (r) shown.

**Figure S6.**
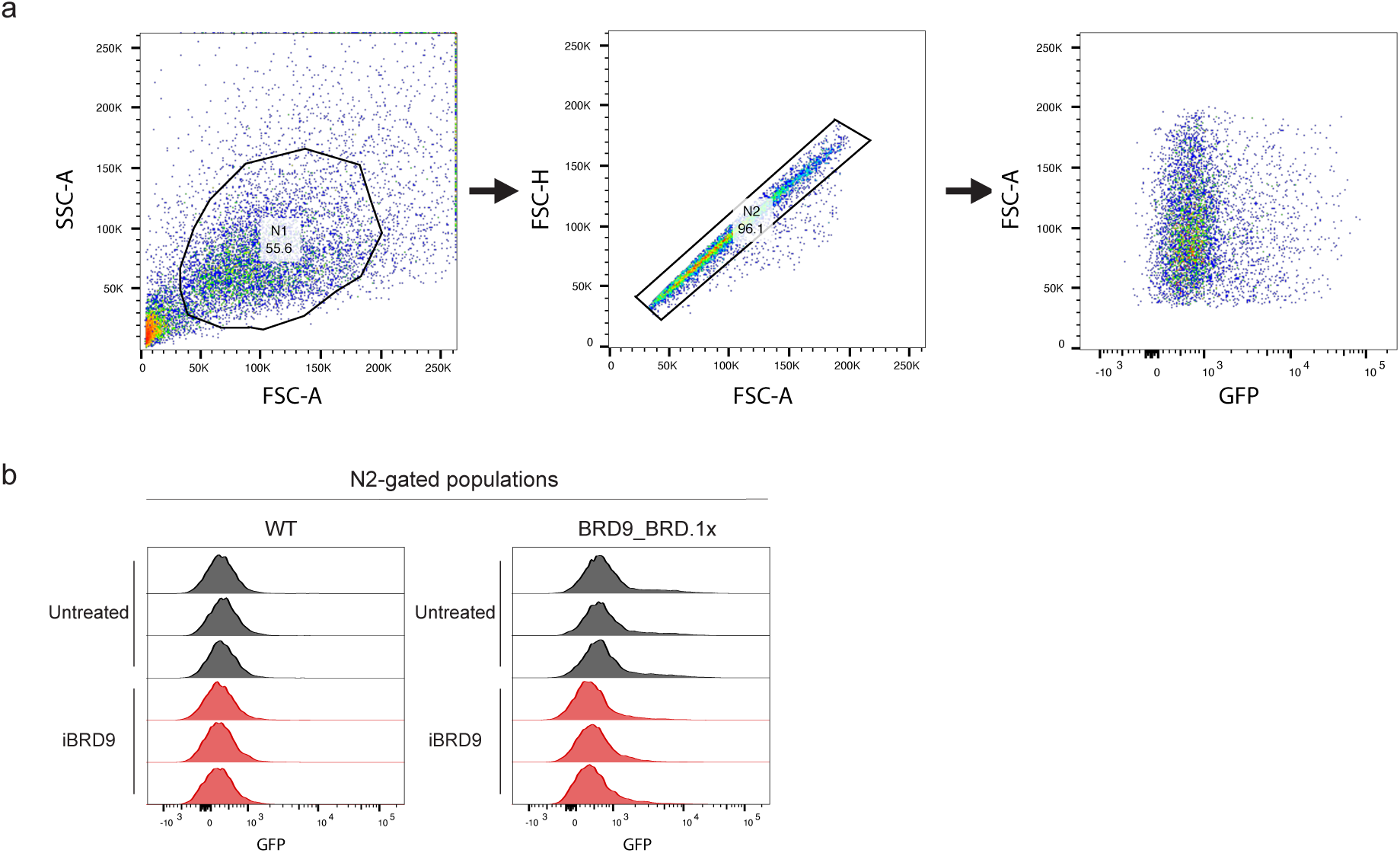
**(A)** Gating strategy for nuclear retention flow cytometry depicted in Figure 5. Individual mESC nuclei were initially gated in region p1. Single nuclei were subsequently gated in region p2 to exclude doublets and debris. eGFP fluorescence signal was collected using the 530/30 nm bandpass filter with 488 nm excitation. **(B)** Nuclear retention flow cytometry analysis of eGFP fluorescence in multiple cell lines after different drug treatments. Three randomly selected replicates per treatment condition are shown for each cell line to show variability amongst replicates. The plots display the mean GFP intensity of nuclsei within the N2 gate, which isolates single nuclei.

